# A species-specific signature residue in the PB2 subunit of the bat influenza virus polymerase restricts viral RNA synthesis

**DOI:** 10.1101/2021.04.12.439585

**Authors:** Saptarshi Banerjee, Aratrika De, Nandita Kedia, Wang Linfa, Arindam Mondal

## Abstract

Bat influenza A viruses (H17N10 and H18N11) are genetically distant from conventional influenza A viruses and replicates poorly in non-bat hosts species. However, the reason behind the lower replication fitness of these viruses are yet to be elucidated. In this work, we have identified species-specific signature residues, present in viral PB2 protein, which is a major determinant of polymerase fitness in human, avian and bat cell lines. Through extensive sequence and structural comparison between the bat and non-bat influenza virus RNA polymerases, we have identified a previously uncharacterized PB2-282 residue, which is serine in bat virus PB2 protein but harbours highly conserved glutamic acid in conventional influenza A viruses. Introduction of these bat specific signatures in the polymerase of a human adapted strain of influenza A/H1N1 virus drastically reduces its polymerase activity and replication efficiency in cell lines of human, bat and canine origin. In contrast, introduction of the human specific signatures in bat virus PB2 (H17N10), significantly enhances its function in the context of a chimeric RNA polymerase. Interestingly, the PB2-282 resides within an evolutionary conserved “S-E-S” motif present across different genera of influenza viruses but is replaced with a “S-S-T” motif in bat influenza viruses, indicating that this E to S transition may serve as a species-specific adaptation signature that modulates the activity of bat virus polymerase in other host species.

**Importance:** Recent isolation of influenza A like viruses (H17N10 and H18N11) in bats raised concerns about their potential of zoonotic transmission in human. Here we present species-specific signature residues present in the bat influenza virus polymerase, which may act as critical modulators of bat virus propagation in non-bat host species. We utilize bioinformatics based comparative analysis followed by functional screening in order to identify the PB2-282^nd^ position, which harbors a highly conserved glutamic acid in conventional influenza A viruses, but contains an unusual serine in case of bat influenza viruses. Human adapted polymerase, harboring bat specific signature (PB2-S282) performs poorly, while bat PB2 protein harboring human specific signature (PB2-E282) shows increased fitness in human cells. Together, our data identifies novel species-specific signatures present within the influenza virus polymerase that may serve as a key factor in the adaptation of influenza viruses from bat to non-bat host species and vice versa.

## Introduction

Influenza A viruses are segmented negative sense RNA viruses that cause respiratory infections in humans and wide range of other animals including, but not limited to, birds, pigs, dogs, cats, horses and seals (1). Wild aquatic birds are the natural reservoirs for these viruses, which through continuous adaptation in their genetic architecture acquires ability to infect new host species. Such adaptation often time results in a new event of zoonotic infection leading to the expansion in the repository of human infecting influenza A viruses (1). For successful zoonoses, influenza viruses have to conquer two barriers imposed by the host; first, successful recognition of the cell surface receptors by viral HA protein to mediate entry (2), and second, species specific adaptation of the viral RNA dependent RNA polymerase (RdRp)(3), majorly governed by its PB2 subunit. The PB2 protein of avian viruses contains a glutamic acid residue at the 627^th^ position while for the human adapted viruses, the same position harbors a lysine. The polymerase containing avian type signature residue in PB2 is fully functional in avian cells, but shows severe attenuation in human cells (4, 5). Other mutations within the 627 domain of PB2 have also been reported to act as species specific adaptation sites that helps the avian viruses to establish successful infection in humans/ mammals (6).

During 2012 and 2013, genetic traces of two new lineages of influenza A viruses were isolated from the little yellow-shouldered bat (*Sturnira lilium*), in Guatemala, and the flat-faced fruit-eating bat (*Artibeus planirostris*), in Peru respectively (7, 8). According to the phylogenetic analyses, these two viruses shows significant divergence in their genetic architecture from classical influenza A viruses, classified as two new subtypes, the H17N10 and H18N11. The H17 and H18 proteins do not recognize the canonical sialic acid receptors (8, 9), while the N10 and N11 lacks sialidase activity (10), suggesting that bat influenza viruses utilize distinct mechanism for entry into the host cells as compared to the classical influenza A viruses. Recent advancements with bat influenza viruses further substantiated this notion through identification of MHC class II (MHCII) as bonafied receptors for bat influenza viruses (11, 12). This also has raised the concerns of epizootic and zoonotic transmission of these viruses, given the constitutive expression of MHCII molecules in wide variety of tissues in human, pigs, chicken and wide range of other animals (11).

Polymerases from human or avian origin influenza viruses have shown to quickly adapt in bat cell lines (13). However, adaptability of the bat influenza virus polymerase in avian, human, or in other mammalian hosts has yet to be extensively investigated. Chimeric bat viruses that express surface antigens (HA and NA) of classical influenza viruses (influenza A/H1N1/PR8 or A/H7N7/SC35M) but internal proteins from the bat influenza virus (H17N10/Guatemala/164/2009) replicates poorly in cell culture and in infected animals compared to the H1N1 (PR8) or H7N7 (SC35M) viruses (14, 15). Additionally, recombinant bat virus polymerase reconstituted in human lung epithelial cells shows limited activity in a reporter-based polymerase activity assay (7). Together, these evidences suggest that bat influenza virus polymerases are severely restricted in non-bat hosts, possibly due to the higher genetic diversity of the internal genes (16) that accommodate unique signature residues in the individual subunit proteins, PB2, PB1 and PA. In this regard it should be noted that the bat virus PB2 protein harbors an unusual serine residue at the 627^th^ position instead of highly conserved glutamic acid or lysine as in case of avian or human adapted viruses respectively.

In this work, we characterized unique molecular signatures present in the bat influenza virus PB2 protein that could be responsible for the compromised fitness of the polymerase in non-bat hosts. Considering the PB2-S627 as signature for the bat virus polymerase, we tried to identify more such positions in the PB2 protein, which harbors either serine or threonine in the bat virus but contains highly conserved glutamic acid or aspartic acid in the non-bat IAV’s. We have identified the Serine at 282 position as a key signature residue, responsible for restricted activity of the bat virus polymerase, in comparison to the human or avian adapted polymerases harboring a glutamic acid residue at the same position. Replacing the glutamic acid either with alanine, lysine or with bat specific serine residue severely affects the RNA synthesis activity of the human virus polymerase when reconstituted through transient transfection. Furthermore, recombinant human viruses with alanine or bat specific serine at 282 position is severely restricted in cell lines derived from human, bat and canine, establishing the importance of the glutamic acid at this position. This is further supported by the fact that partially humanized H17N10 PB2, harboring glutamic acid at 282 position boosts polymerase activity several folds compared to its wildtype version harboring serine at the same position. Together, our work identified a key molecular signature in bat influenza virus PB2 protein which is responsible for its restricted activity in non bat host species and may serve as a potential site of adaptation in order to acquire improve fitness in other hosts including human.

## Results

### Identification of the bat specific signatures responsible for the restricted activity of the polymerase in non-bat host

In order to identify the species-specific signature residues in the bat virus polymerase we focused upon the PB2 subunit due to its critical role in determining the species specificity of influenza A viruses (4, 5). Throughout this study the influenza A/H1N1/WSN/1933 strain was used as a model for the human adapted virus while the influenza A/H17N10/Guatemala/60 strain was used as a representative for the bat virus. First, we replaced the human-specific lysine at the 627^th^ position with the bat-specific serine residue in order to ascertain its impact upon the activity of the human virus polymerase in human (HEK293T) and avian (DF1) cells. Luciferase based reporter RNPs were reconstituted through transient transfection of a genome sense reporter RNA template, polymerase subunits (PB1, PB2 and PA) and NP expressing plasmids derived from the A/H1N1/WSN/1933 strain as described previously (Mondal et al, 2015). Different variants of PB2 harboring the human signature lysine (K), avian signature glutamic acid (E) or bat signature serine (S) residues at the 627^th^ position were used to reconstitute the polymerase (Fig1.A). The polymerase with PB2-627K shows high reporter activity in both the cell lines while the polymerase with PB2-627E was selectively attenuated in human cells as expected (4, 5, 18), hence validating the efficiency of our luciferase-based assay system in determining the species-specific fitness of the polymerase (Fig1.B). Interestingly, the polymerase with PB2-627S still remained attenuated, although to a lesser extent than PB2-627E, in human cells but was fully functional in chicken cells (Fig1.B). Different PB2 variants shows comparable expression as evidenced from the western blot analysis (Fig1.B). These data suggest that the PB2-627S may serve as one of the key determinants for species specific restriction of the bat virus polymerase in non-bat host species, specifically in human.

**Figure 1.**
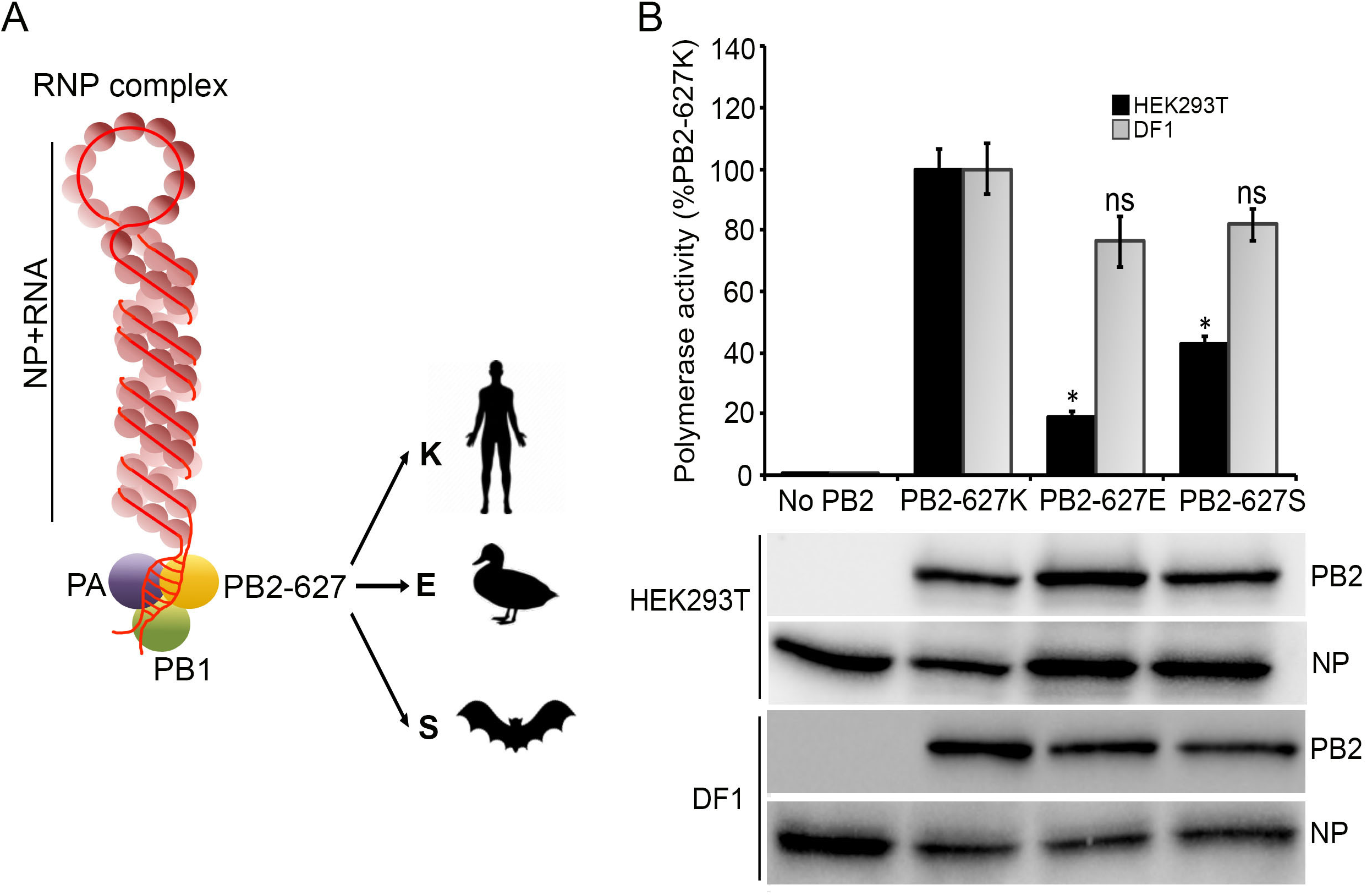
Bat specific serine at PB2-627 position restricts the polymerase in human but not in avian cells. **A**. Influenza A virus RNPs are the major determinant of its host specific replication fitness. The 627^th^ amino acid of the PB2 subunit of polymerase harbors a lysine (K) or a glutamic acid (E) residues for human or avian adapted viruses respectively. Bat influenza viruses harbors a serine (S) at the same position. **B**. Luciferase based reporter assay was performed to assess the polymerase function in human and chicken cells. Viral RNP, with the genetic background of A/WSN/1933, was reconstituted in HEK293T or DF1 cells either with wildtype PB2 containing a lysine, or mutant PB2 containing glutamic acid or serine residue at 627th position. n=3±standard deviation. *p<0.05 one-way ANOVA when compared to PB2-627K.

Encouraged by this finding, we focused our study towards identification of additional signature residues, present in the bat influenza virus PB2 protein, which may regulate the activity of viral RNA polymerase in a species-specific manner. Using the PB2-627S as a reference, we tried to identify specific positions throughout the primary sequence of PB2 protein, which harbors either serine or threonine in the bat virus polymerase but represents highly conserved glutamic acid or aspartic acid in the conventional influenza viruses of either human or avian origin. Through extensive alignment of the bat and not bat influenza A virus PB2 sequences we have been able to identify total seven such amino acid residues (60, 282, 390, 472, 671, 678, 681), which were then further screened based upon their solvent accessibility in the atomic structure of the polymerase (19, 20). Finally, we focused upon five amino acid residues, 282, 390, 472, 678, 681 that are completely surface exposed (Fig2.A, table 1) in the heterotrimeric polymerase (in analogy with the PB2-627 position), harbor serine in bat virus PB2 protein but are occupied by highly conserved aspartic acid (390, 678) or glutamic acid (282, 472, 681) in the non-bat IAV’s (Table1 and Fig1A). Residue number 60 and 671 were not included in the study as they are not completely surface exposed in the heterotrimeric polymerase rather reside within the PB2-PB1 and PB2-PB1-PA interaction interface respectively. We speculated that some of these residues may serve as a determining factor in regulating the bat virus polymerase activity in bat and (or) non-bat host species.

**Figure 2.**
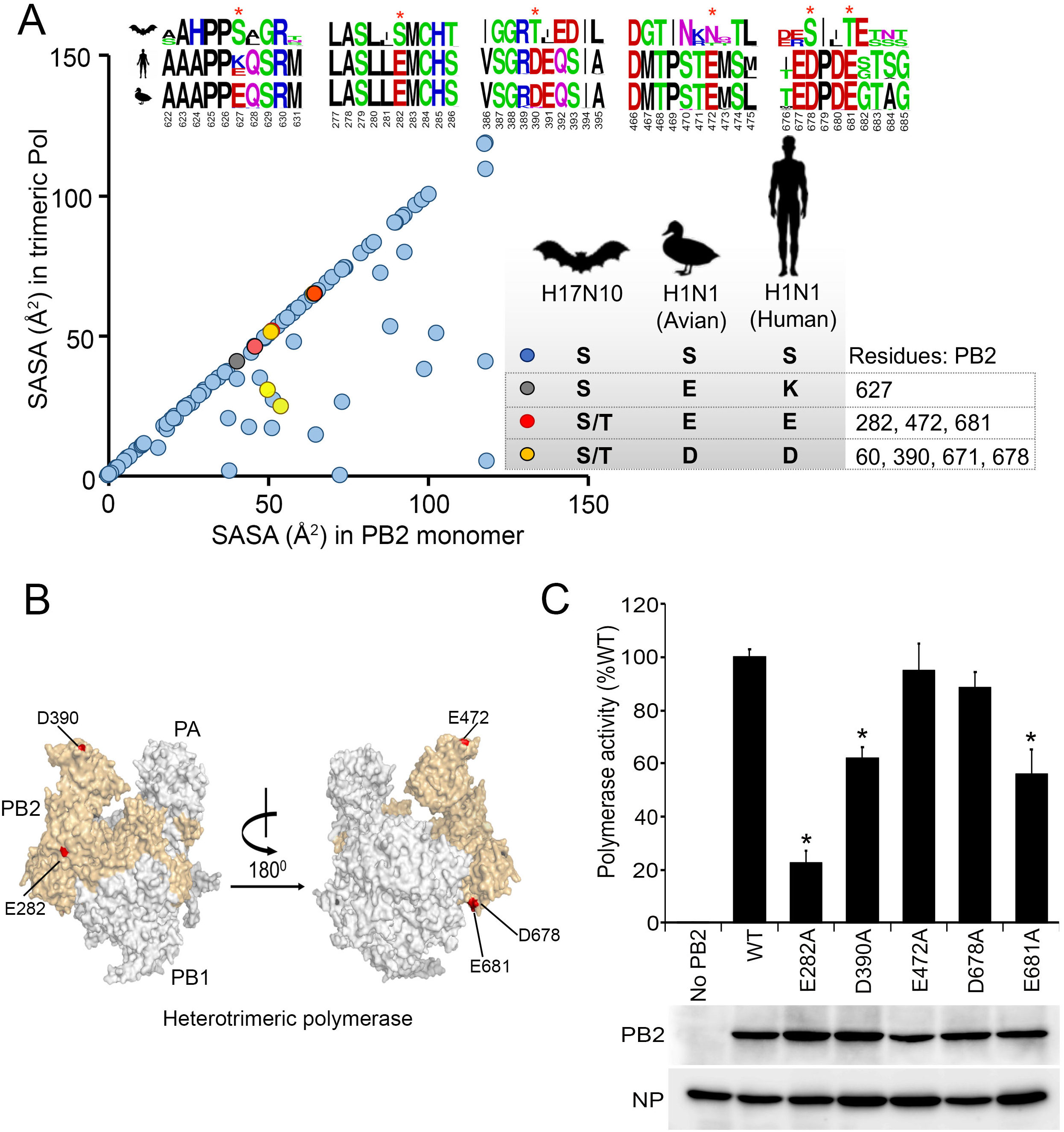
Identification of species-specific signature residues in PB2 subunit of bat influenza virus polymerase. **A**. Residue wise solvent accessible surface area (SASA in Å^2^) of H17N10 monomeric PB2 and in trimeric polymerase was calculated. Serine and threonine residues were plotted as blue dots on the basis of their exposed surface area in monomeric and trimeric form. Serine or threonine residues in bat virus PB2 protein, which are a glutamic acid or aspartic acid residue in conventional Influenza viruses are marked as red and yellow dots respectively. Logo-plots of the shortlisted amino acid residues (marked with Asterix) which represent highly conserved glutamic /aspartic acid in the PB2 protein of influenza A viruses of human (sequences analyzed: 36243) or avian (sequences analyzed: 19762) origin but harbors serine in the PB2 of bat viruses (sequences analyzed: 7). **B**. Spatial organization of the surface exposed glutamic /aspartic acid residues in the heterotrimeric polymerase structure of a human adapted influenza A virus (A/NT/60/1968,PDB ID:6RR7) that are shortlisted for functional screening. **C** Luciferase based reporter assay was performed to assess the polymerase function of the recombinant RNA dependent RNA polymerase with wild type or alanine substitution mutants of PB2 proteins using the genetic background of A/WSN/1933 strain. n=3±standard deviation. *p<0.05 one-way ANOVA when compared to wildtype PB2.

**Table 1:** **Analysis of surface exposed serine and threonine residues in H17N10 and H3N2 PB2 from the crystal structure of RNA dependent RNA polymerase (4WSB.PDB &6RR7.PDB)**.

**(4WSB.PDB & 6RR7.PDB).**
| Criteria | Number | Position |
| --- | --- | --- |
| Total number of surface exposed serine and threonine residues in H17N10-PB2 (4WSB.PDB) | Serine (34),<br>Threonine (34) | **Serine-**<br>14,61,92,156,178,179,225,226,279,282,288,<br>290,291,300,322,324,366,371,442,443,481,5<br>14,533,534,544,593,622,674,678,683,684,68<br>5,688,741<br>**Threonine-**<br>76,79,129,155,184,186,224,227,235,245,286<br>,287,303,340,346,351,353,355,390,400,444,<br>451,468,472,549,562,566,569,582,596,598,6<br>09,631,681 |
| Total number of surface exposed serine and threonine residues in H3N2-PB2 (6RR7.PDB) | Serine (20)<br>Threonine (20) | **Serine –**<br>14,92,179,225,226,279,286,322,324,481,514<br>,533,534,544,593,684,688,79,155,286,582<br>**Threonine-**<br>178,371,674,683,76,129,184,186,224,235,24<br>5,287,303,346,351,468,549,569,598,609 |
| Surface exposed<br>Serine and<br>Threonine<br>residue in<br>H17N10-PB2<br>changed to other<br>residue in<br>classical<br>influenza virus<br>PB2 | Serine (17)<br>Threonine (18) | **Serine-**<br>61,156,178,282,288,290,291,300,366,371,44<br>2,443,622,674,678,683,685<br>**Threonine-**<br>76,155,227,286,340,353,355,390,400,444,45<br>1,472,562,566,582,596,631,681 |
| Surface exposed<br>Serine and<br>Threonine<br>residue of<br>H17N10-PB2<br>changed to<br>aspartic and<br>glutamic residue<br>in classical<br>influenza virus<br>PB2 | Aspartic acid<br>(2)<br>Glutamic acid<br>(3) | **Glutamic acid-**<br>282,472,681<br>**Aspartic acid-**<br>390,678 |

In order to test this, we first investigated the functional significance of the identified amino acid residues in the context of the human adapted polymerase. For this purpose, we have substituted each of the aspartic acid or glutamic acid residues with alanine in order to generate a panel of the mutant PB2 proteins. Luciferase based reporter RNPs were reconstituted either with wild type or with mutant PB2 proteins along with other RNP components as described earlier. Polymerase reconstituted with the wildtype PB2 protein showed high level of reporter activity in HEK293T cells while mutant PB2 proteins supported polymerase activity to various extents (Fig 2.B). The PB2 D472A and E678A mutants supported polymerase activity comparable to the wild type protein suggesting that these residues are not critical for viral RNA synthesis. The D390A and E681A mutants showed only modest defects reducing polymerase activity only by 40%. Interestingly, the E282A mutation severely attenuated the polymerase, supporting only 20% reporter activity compared to the wildtype PB2. These data suggest that the glutamic acid at 282 position of PB2 is critical in supporting activity of the human adapted polymerase in human cells. Western blot analysis of the E282A and other mutant PB2 proteins showed expression levels comparable to the wild type protein, hence confirming that the defect in RNA synthesis results from the suboptimal activity of these proteins and not from their altered abundance in cells.

### Recombinant influenza A/H1N1/WSN viruses harboring bat specific signature residues at the 627 and 282 position of PB2 shows reduced fitness in bat and not bat host species

Highly conserved PB2-K627 and PB2-E282 residues are functionally important in the context of human virus polymerase suggesting that the bat specific serine residues at corresponding positions may alter the polymerase activity and hence interfere with virus propagation in a species-specific manner. Hence, we investigated the impact of the bat specific serine residues at the 627^th^ and 282^nd^ positions of PB2 in the context of the propagation of human adapted viruses. The genetic background of the influenza A/H1N1/WSN/1933 strains was used and K627S, E282A and E282S mutations were introduced in the PB2 open reading frame in order to generate recombinant viruses using the plasmid based reverse genetics as described earlier(18). Initially rescued viruses were amplified in MDCK cells for three consecutive passages, then titrated using plaque assay and finally confirmed the presence of the mutations in the PB2 ORF through sanger sequencing. While the PB2-627S virus shows plaque sizes comparable to the wild type virus, PB2-282A and PB2-282S viruses showed significantly smaller plaque sizes indicating possible defect in the replication of the mutant viruses (Fig 3A). Interestingly, the E282S mutant virus incorporated a conservative leucine to valine replacement at the adjacent 281 position which occurred consistently for the viruses rescued in two independent experiments. The reverse genetics plasmid harboring the PB2 open reading frame showed no such change in its nucleotide sequence suggesting that the virus spontaneously incorporated L281V mutation in its genomic RNA sequence. It should be noted that the E282 residue is conserved not only for influenza A, but also for influenza B and C viruses, where this glutamic acid residue is preceded by the highly conserved valine as observed in the case of our E282S/L281V mutant virus. Hence, we moved forward with the mutant viruses to investigate their ability to propagate in cell lines from various host species. For simplicity, we designate the E282S/L281V mutant virus as E282S virus for the rest of the manuscript.

**Figure 3.**
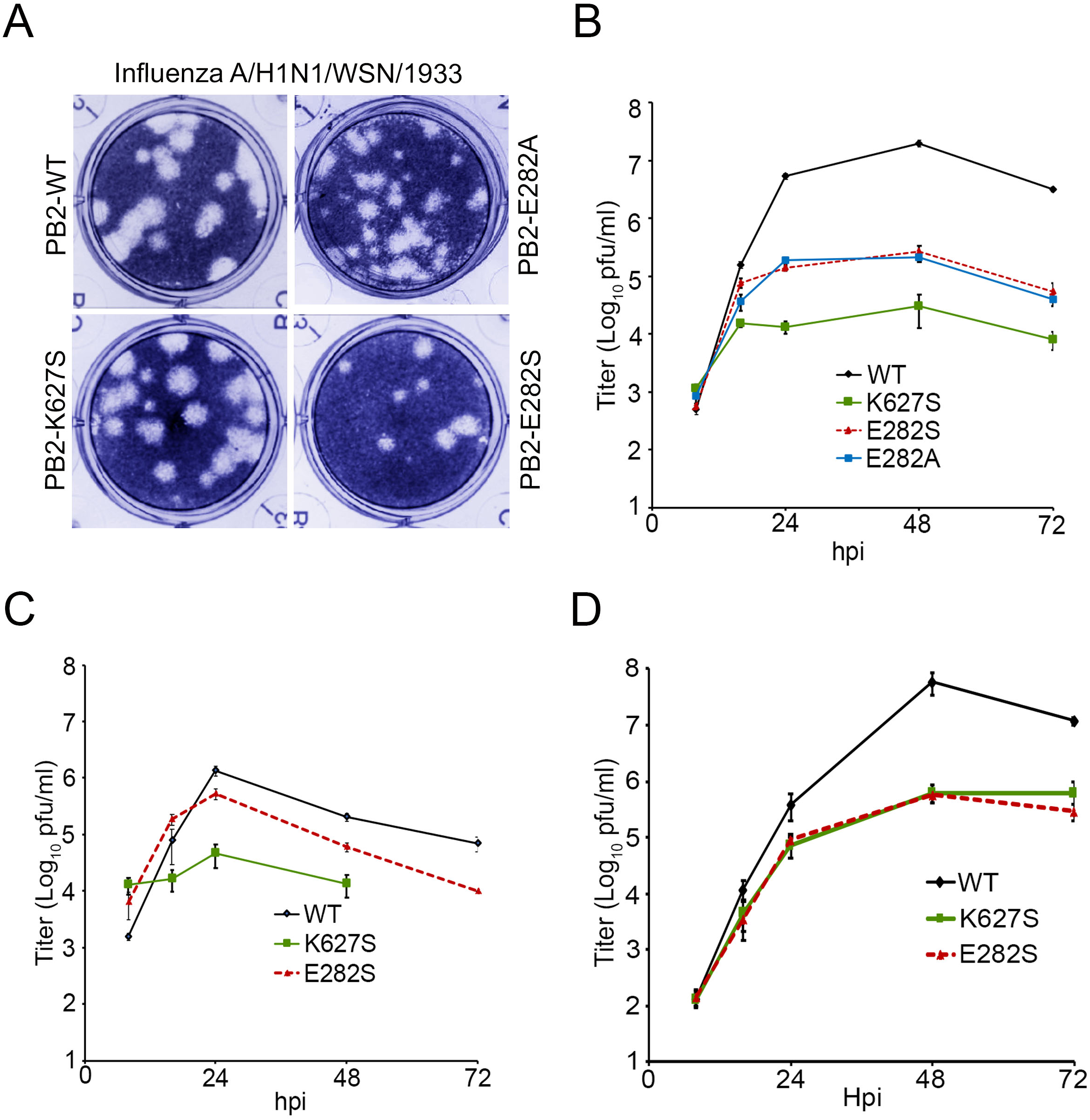
Conventional influenza A viruses, harboring bat specific signature serine residues are defective in bat and not-bat host species. Recombinant influenza A/H1N1/WSN/1933 was generated contains a serine or alanine residue at 282^nd^ position and serine at 627^th^ position of PB2. **A**. Plaque morphology of recombinant viruses contain the mutation. **B**. Madin Darby canine kidney, MDCK, **C**. Human lung epithelial carcinoma, A549 and **D**. *Pteropus Alecto* kidney, PaKi cells were infected with recombinant viruses at an MOI of 0.01 in three biological replicates. Supernatants were collected at 8,16,24,48 and 72 hours post infection. Viral titer was calculated for each time points by performing plaque assay.

First, we have evaluated the fitness of the mutant PB2 viruses by performing multicycle replication kinetics in Madin-Darby Canine Kidney (MDCK) cells. The wildtype virus replicated to high levels reaching titers more than 10^7^ by 48 hours post infection, while the PB2-K627S virus was severely attenuated showing up to 1000-fold reduction in viral titer throughout the time course (Fig3.A). This clearly reflects that introduction of the bat specific residue in the human polymerase severely restricts its activity in cells of mammalian origin, which is consistent of the results demonstrated with others strains of influenza A/H1N1 virus (21). Introduction of either alanine or the bat specific serine residue at the 282-position showed similar attenuation with a 100-fold decrease in the titer with respect to wildtype virus (Fig3.A). Interestingly, the PB2-E282S virus showed higher replication fitness compared to the PB2-K627S virus in MDCK cells. To have a better insight into the specie-specific fitness of human influenza virus that could be modulated by the bat specific serine residues at the PB2-627 and PB2-282 positions, we next evaluated the replicability of the mutant viruses in cell lines of human and bat origin. We have used human lung epithelial carcinoma cells (A549) and *Pteropus alecto* kidney (PaKi) cells as model cell lines of the respective host species. As described earlier, the PaKi cells support influenza virus infection and re-assortment (22), which makes them suitable for monitoring replication of the human viruses harboring bat specific signatures in their replication machinery. In fact, both wildtype and mutant viruses replicated to higher extents in PaKi cells compared to the A549 or MDCK cells showing high susceptibility and permissiveness of these cells towards H1N1 virus infection (Fig3.C). In A549 cells, replication efficiency of the wild type and mutant viruses showed trends similar to what was observed in MDCK cells. The PB2-K627S showed up to 100-fold attenuation followed by the PB2-E282S showing around 10-fold reduction in viral titers as compared to the wild type virus (Fig3.B). Interestingly, both the mutant viruses showed similar replication fitness in PaKi cells, with more than 100-fold reduction in virus titer as compared to the wild type one (Fig3.C). Together our data suggest introduction of the bat specific serine residues at the critical 627 and 282 positions of PB2 can significantly restrict the replication of the prototypic WSN strain of human influenza virus in both bat and non-bat host species. Based upon the sequence and structural conservation of the above-mentioned residues, it is likely that similar phenotypes might be observed for other conventional influenza viruses as well.

### A highly conserved “S_279_-E_282_-S_286_” motif within the mid-link regions of the PB2 protein is crucial for optimum fitness of the polymerase in human and avian cells

The E282 resides within the “mid-link” region of PB2, which connects the N terminal and cap binding domains. As revealed by different atomic structures of the polymerase, the “mid link” region along with the “627-linker”, together forms the stock upon which the cap binding domain and the 627 domain can undergo structural reorganization in order to transition between the transcriptionally active and replicative forms (19, 20). The mid link region consists of four α helices (α14-α17), two of which face towards the core of the polymerase, while the other two are solvent exposed. The E282 resides on the second turn of the α15 helix flanked by two other serine residues in the adjacent turns, all three facing towards the solvent (Fig4.A). Due to this structural feature the three residues, S279, E282 and S286, constitute a unique motif, which is completely conserved across the PB2 sequences derived from the conventional influenza A viruses. Furthermore, extensive structural and sequence alignment studies revealed that this S-E-S motif is also conserved in influenza C viruses and partially conserved in influenza B and D viruses (Fig4.A, B). Interestingly, the bat influenza A virus PB2 harbors a serine at the 282 position and a threonine at 286, which results in an altered S_279_S_282_T_286_ motif with the same structural feature (Fig4.B).

**Figure 4.**
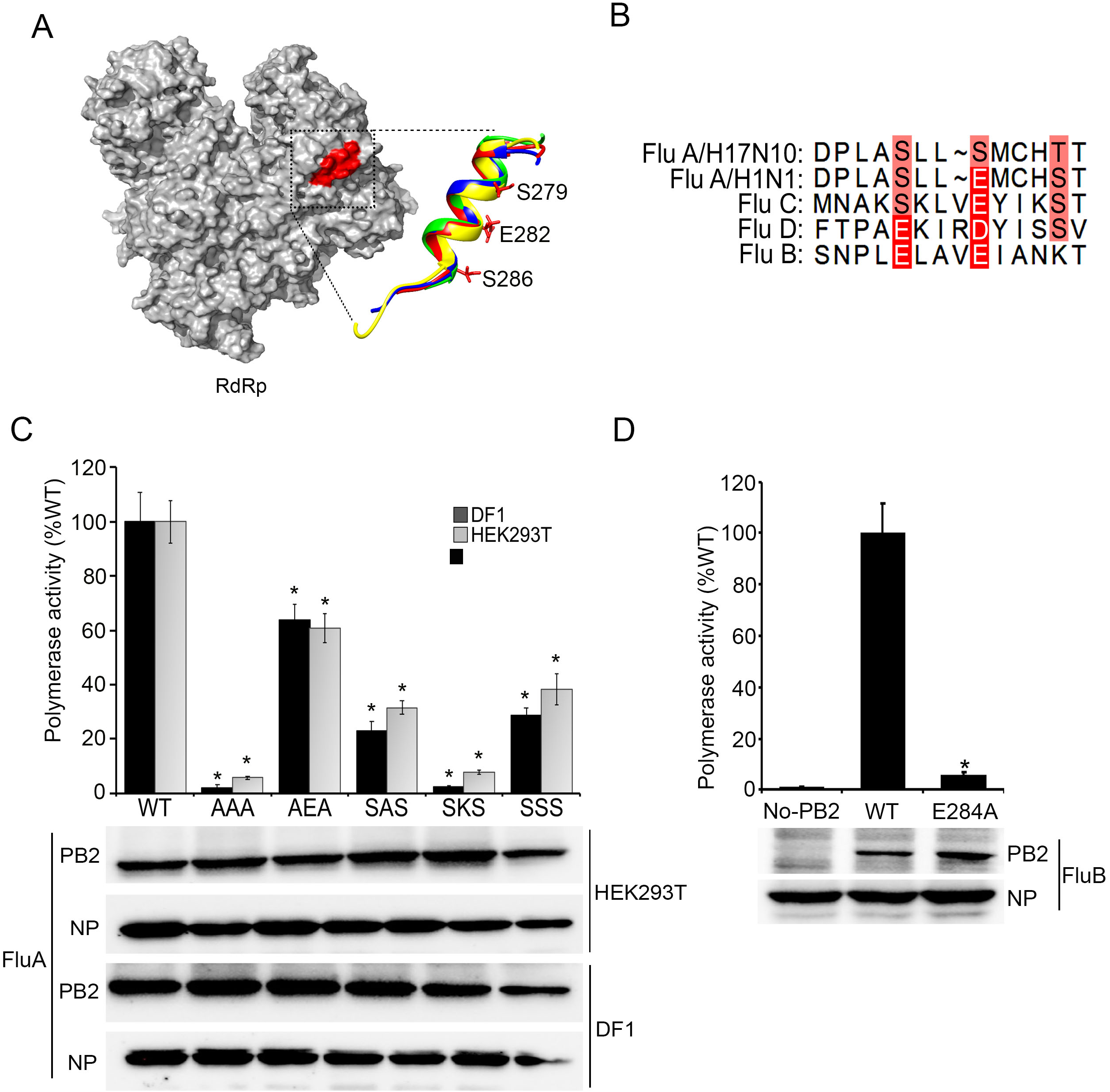
A surface exposed, highly conserved “S-E-S” motif is crucial for the activity of viral RNA polymerase. **A**. PB2 282^nd^ residue is present in the α15 helix (highlighted in red color in the surface representation of the polymerase) of the mid-link region, which is completely surface exposed and structurally conserved in influenza A (red), bat infecting influenza (blue), influenza B (yellow) and influenza C (green) viruses. The amino acid residues denoted are of influenza A. **B**. Multiple sequence alignment of Influenza A/H17N10(A/H17N10/Guetamala/060), Influenza A /H1N1 (A/H1N1/WSN/1933), influenza C (C/Aichi/1/81), influenza D (D/Quebec/1M-H/2019) and influenza B (B/Brisbane/163/2008) showing the variable degrees of conservation of the S-E-S motif. **C**. Luciferase based reporter assay was performed to assess the RNA dependent RNA polymerase function (using the genetic background of influenza A/H1N1/WSN/1933 strain) in human and chicken cells reconstituted by wild type or mutant PB2 protein. n=3±standard deviation. *p<0.05 one way ANOVA when compared to wildtype PB2. **D**. Luciferase based reporter assay was performed to assess the polymerase function of Influenza B/Brisbane/2008 RNA dependent RNA polymerase containing wild type or mutant PB2 protein. n=3±standard deviation.*p<0.05 one way ANOVA when compared to wildtype PB2.

Based upon the importance of the E282 position in supporting viral RNA synthesis and virus replication and the high conservation of the “S-E-S” motif across different influenza virus PB2 protein, we next evaluated the importance of this motif in supporting polymerase activity in HEK293T and DF1 cells. For this purpose, we generated a series of mutant PB2 proteins where individual amino acids of the S-E-S motif were substituted either with alanine, serine or lysine and tested their ability to support viral RNA synthesis using the reporter-based polymerase activity assay (Fig4.C). Substitution of both of the peripheral serine residues with alanine resulted in 40% decrease in reporter activity, while substitution of the central glutamic acid showed around 70-80% decrease (Fig4.C), as also evidenced earlier (Fig2.B). A triple alanine mutant, substituting the entire S-E-S motif resulted in complete abrogation of RNA synthesis, pointing towards the importance of the entire motif in supporting the activity of viral RNA polymerase. It is interesting to note that introduction of bat specific serine residue at 282 position and hence reconstituting the bat like “SSS” motif resulted in around 60-70% decrease in polymerase activity, further emphasizing the importance of the negatively charged glutamic residue at the center of the motif (Fig4.C). This is also substantiated by the fact that substitution of the negatively charged glutamic acid with positively charged lysine results in complete abrogation of the polymerase activity. All of the mutant proteins show expression and stability comparable with the wild type PB2 protein as evidenced from the western blot analysis (Fig4.C). Together our data revealed the existence of a highly conserved “S-E-S” motif present in the mid link region of influenza A virus PB2 protein, which is crucial in supporting the polymerase activity both in avian and human cell lines.

Extensive structural and sequence comparison between influenza A and B virus PB2 proteins suggest that the B-PB2 protein lacks the canonical “S_279_-E_282_-S_286_” motif, in spite of having significant sequence similarity within the mid-link region (Fig4.B). Instead, it contains two glutamic acid residues in the two subsequent turns of the alpha helix, hence constituting the “E_280_-E_284_” motif where the E-284 of B-PB2 perfectly aligns with the E282 of A-PB2 protein (Fig4.B). To investigate whether the conserved glutamic acid at 284 is important for influenza B polymerase activity, we have developed a firefly luciferase-based influenza B polymerase activity assay using the genetic background of B/Brisbane/60/2008 strain. Influenza B polymerase with wild type PB2 supported high levels of reporter activity as shown in figure 4D. Interestingly, introduction of the E284A mutation in B-PB2 reduced the reporter activity by 84%, suggesting indispensable role of this glutamic acid in supporting polymerase activity. This data further substantiates the importance of this highly conserved glutamic acid residue in the PB2 protein of influenza viruses across different genera. This also points towards the fact that absence of such an important molecular signature may significantly impact efficiency of the bat influenza virus PB2 protein, thereby restricting the overall activity of the bat virus polymerase in comparison to the conventional influenza A or influenza B viruses.

### Alteration of the “SES” motif specifically restricts the RNA synthesis ability of the polymerase

Importance of the PB2 “S-E-S” motif, specifically the E282 residue, in supporting influenza A virus polymerase activity motivated a detailed investigation of its precise role in viral RNA synthesis. Influenza viruses assemble their RNA synthesis machinery in the form of viral ribonucleoprotein complexes (RNPs), where the heterotrimeric RNA polymerase (PB1, PB2 and PA) recruits multiple copies of NP on the nascent RNA strand during its synthesis, resulting in the formation of a macromolecular RNA-protein complex of megadalton range. Hence, fruitful assembly of viral RNPs are prerequisite for subsequent rounds of RNA synthesis. To investigate the molecular mechanism by which mutations introduced in the S-E-S motif altered viral RNA synthesis, we assessed their ability to support viral RNP assembly using RNP reconstitution. Authentic viral RNPs are reconstituted in HEK293T cells by expressing the negative sense genomic RNA template, NP, PB1, PA and wild type or mutant variants of Flag tagged PB2 proteins. The efficiency of RNP formation was determined by immunoprecipitating the viral polymerase via PB2-FLAG and detecting co-precipitated NP as a part of RNP complex by western blot analysis. A strong signal of NP reflects co-precipitation of large number of NP molecules as a part of RNP complex, while faint signal represents limited coprecipitation as a result of direct PB2-NP interaction. As evidenced in figure5.A, wild type PB2 efficiently co-purified large portions of NP indicating efficient RNP formation. The AAA and the SKS mutants completely abolished the RNP formation while the SAS mutant also showed a drastic reduction. AEA and the SSS mutants showed and intermediate phenotype, showing higher levels of RNP formation than SAS mutant but lower than the wild type protein. Together, the ability of the PB2 mutants to support RNP assembly perfectly corroborates with their ability to support viral RNA synthesis as evidenced in the reporter activity assay.

**Figure 5.**
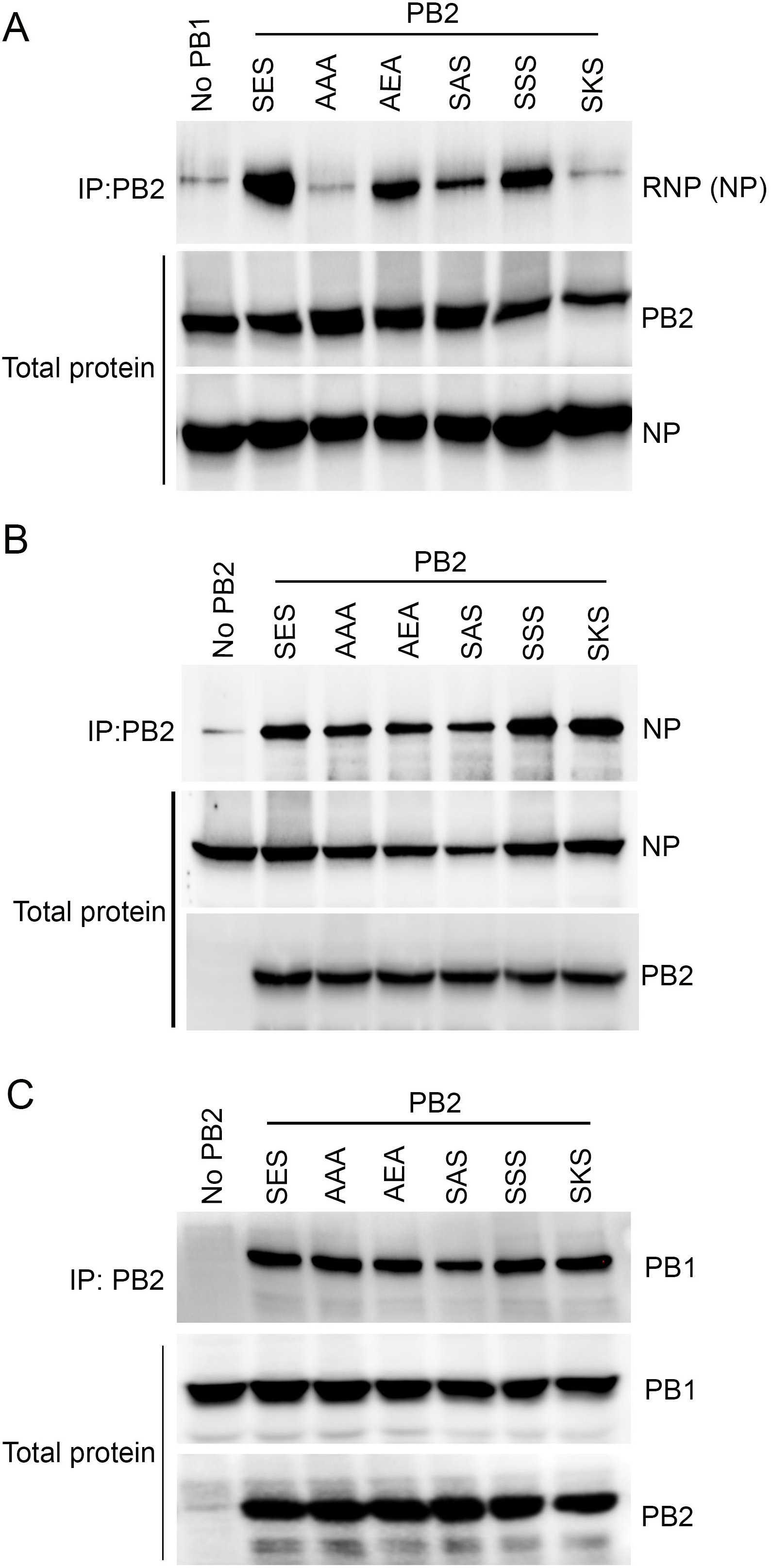
Alteration of the “SES” motif affects RNP reconstitution without impacting interaction with other viral proteins, NP and PB1. **A**. RNP reconstitution was performed by transfecting NP, PB1, PA, PB2 protein and vRNA expression plasmid in HEK293T cells. 48 hours post transfection cells were lysed and PB2 protein was immunoprecipitated and blotted for co-precipitated NP. **B**. PB2 and NP interaction was checked by transfecting NP expressing plasmid along with wild type or mutant PB2 expressing plasmids in HEK293T cells. 48 hours post transfection cells were lysed and PB2 was immunoprecipitated and blotted for co-precipitated NP. **C**. PB2 and PB1 interaction was checked by transfecting PB1 expressing plasmid along with wild type or mutant PB2 expressing plasmids in HEK293T cells. 48 hours post transfection cells were lysed and PB2 was immunoprecipitated and blotted for co-precipitated PB1

Fruitful assembly of RNPs relies upon efficient binary interactions between PB2 with PB1 and NP. Hence, we tested the ability of the mutant PB2 proteins to participate in these protein-protein interactions. Flag tagged wild type or mutant PB2 proteins were co-expressed either with NP or with PB1 in HEK293T cells and the lysates containing the binding partners were subjected to PB2-FLAG immunoprecipitation followed by western blotting to detect the coprecipitation. To eliminate any non-specific complex formation with cellular RNA, lysates were treated with high amounts of RNase A before immunoprecipitation. All of the PB2 mutants showed NP and PB1 coprecipitation comparable to the wildtype protein suggesting mutations in the S-E-S motif does not impact the ability of PB2 to interact with other RNP associated proteins (Fig5.B & C). We have also evaluated the nuclear localization ability of PB2 mutants by overexpressing them in A549 cells followed by indirect immunofluorescence assay. PB2 contains nuclear localization signal which directs its importin alpha mediated nuclear import in order to participate in RNP formation (23, 24). All of the mutant proteins showed preferential nuclear accumulation comparable to the wild type PB2, when expressed through transient transfection in HEK293T cells (Fig.6). Based upon these data it can be predicted that neither impaired protein-protein interaction nor alteration of the subcellular localization was the key for the altered RNP formation ability as evidenced in case of the PB2 mutants.

**Figure 6.**
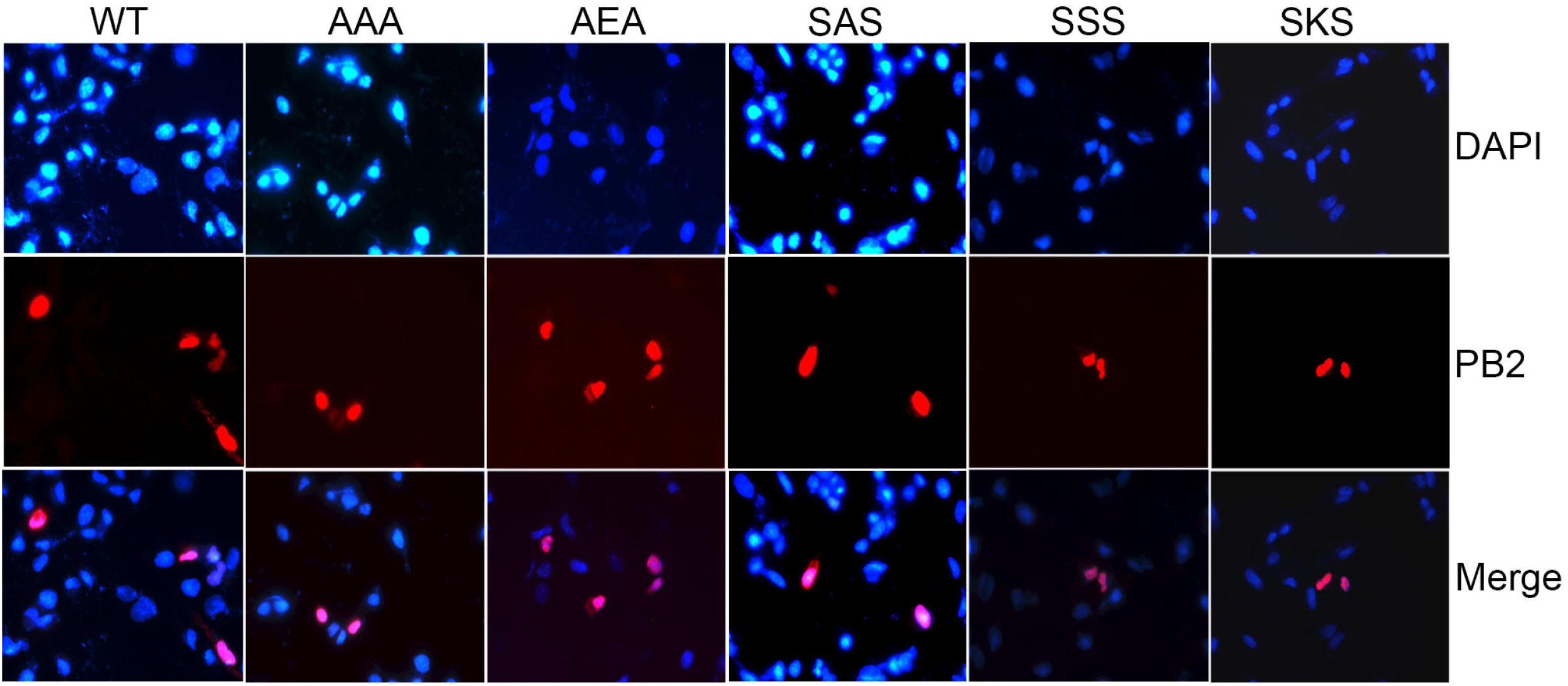
Motif mutants shows similar subcellular localization to the wild type PB2 protein. Plasmids expressing wild type of mutant PB2 proteins were transfected in A549 cells. 24 hours post transfection cells were fixed and parmealized. Cells were first stained with anti-PB2 mouse antibody followed by Alexa fluore 488 anti-mouse rabbit antibody. Nucleus were stained with DAPI. Image were taken in Leica Fluorescence microscope.

Synthesis of viral genomic/ antigenomic RNA and their assembly into progeny viral RNPs takes place in a concomitant fashion. Hence, any defect in viral RNA synthesis would get manifested in the form of defective RNP assembly and vice versa. To test whether PB2 mutants are actually defective in their ability to support viral RNA synthesis, we have performed a mini-vRNA template-based primer extension assay where the RNA synthesis activity of the polymerase could be monitored independent of RNP assembly process. The mini-vRNA templates are short RNAs (76 nucleotide long) containing viral UTRs that could be replicated and transcribed by the polymerase in the absence of NP (25). Hence, any defect in RNA synthesis could solely be attributed to the defect in polymerase activity and not to the defective RNP assembly process. Mini-vRNA template (NP-77) was expressed in HEK293T cells along with the polymerase constituted either with wildtype of mutant PB2 proteins and primer extension assay was performed in order to monitor vRNA (replication) and mRNA (transcription) synthesis in absence of NP. As evidenced clearly, PB2 mutants shows various degrees of defects in vRNA synthesis (Fig.7A & B), which could perfectly be correlated with their ability to support polymerase activity (evidenced in reporter assay, Fig 3.C) and RNP assembly process. Synthesis of mRNA was almost completely abrogated for all of the mutants (only AEA mutant showed minimal activity) (Fig7.A & B), which could be a consequence of the low abundance of the vRNA template due to defect in replication process. This data clearly elucidates an inherent defect in RNA synthesis activity of the polymerase harboring the PB2 mutant proteins with alteration in their SES motif.

**Figure 7.**
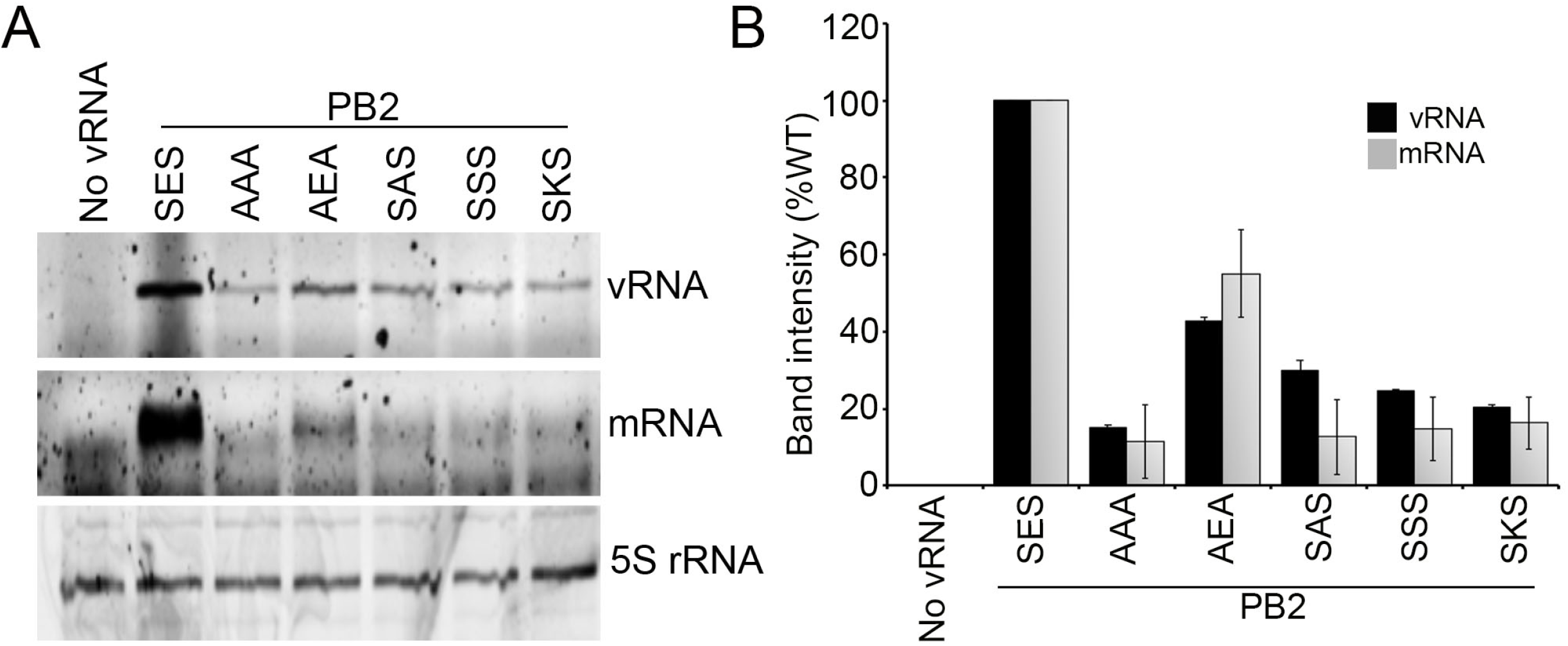
The “SES” motif is critical for supporting RNA synthesis activity of the polymerase. **A**. primer extension was performed with a 77 nucleotide long mini viral RNA template containing 3’ and 5’ UTR of NP gene. Trimeric polymerase was reconstituted in HEK293T cells with wild type or Mutant PB2. Total RNA was extracted by TRIzol method and primer extension performed with fluorescently labelled specific primer for vRNA, mRNA and 5srRNA. Reaction products were analyzed by Urea-PAGE and imaged in Bio-Rad chemidoc. **B**. Densitometric analysis of primer extension experiment from three independent experiments.

### Introduction of human virus specific residues in the bat virus PB2 protein boosts the activity of the chimeric polymerase

Extensive investigation of the molecular mechanism revealed that the highly conserved S-E-S motif, specifically the E282 residue, in the mid-link region of the PB2 is critical in supporting optimum activity of classical influenza A virus polymerase. Alteration of the SES motif via introduction of the bat specific serine residue at 282 position significantly restricts polymerase activity and hence inhibits virus propagation not only in human but also in bat cells (Fig3.C). Similar attenuation was also observed in case of viruses harboring the bat specific serine residue at the 627 position (Fig3.C)(21). Based upon these observations we hypothesize that the serine residues at critical positions of PB2 may serve as the key restriction elements present within the bat virus polymerase that are responsible for the lower replication fitness of bat influenza virus in comparison to the classical influenza A viruses.

To test our hypothesis, we reconstituted a chimeric polymerase composed of the PB1 and PA subunits of the A/H1N1/WSN/1933 strain and the PB2 subunit of the A/H17N10/guatemala/060 strain. Influenza A/H1N1/WSN/1933 virus RNPs were reconstituted either with wild type or chimeric polymerases in HEK293T cells as described earlier (13). As evidenced, introduction of the bat virus PB2 in the human virus RNP severely restricts the polymerase resulting in around two logs decrease in reporter activity (Fig8.A). Although an increasing concentration of the H17N10 PB2 results in a dose dependent increase in reporter activity, the chimeric polymerase still remained attenuated with respect to the wild type, suggesting the severe restriction imposed by the bat PB2 upon the other subunit of the polymerase. This is in corroboration with the results presented by other groups, where alteration of the individual RNP component of human adapted influenza virus with that of the bat influenza virus restricted viral RNA synthesis (Pool et al, 2014; Juozapaitis et al., 2014; Zhou et al., 2014). Subsequently, we have replaced the bat specific serine

**Figure 8.**
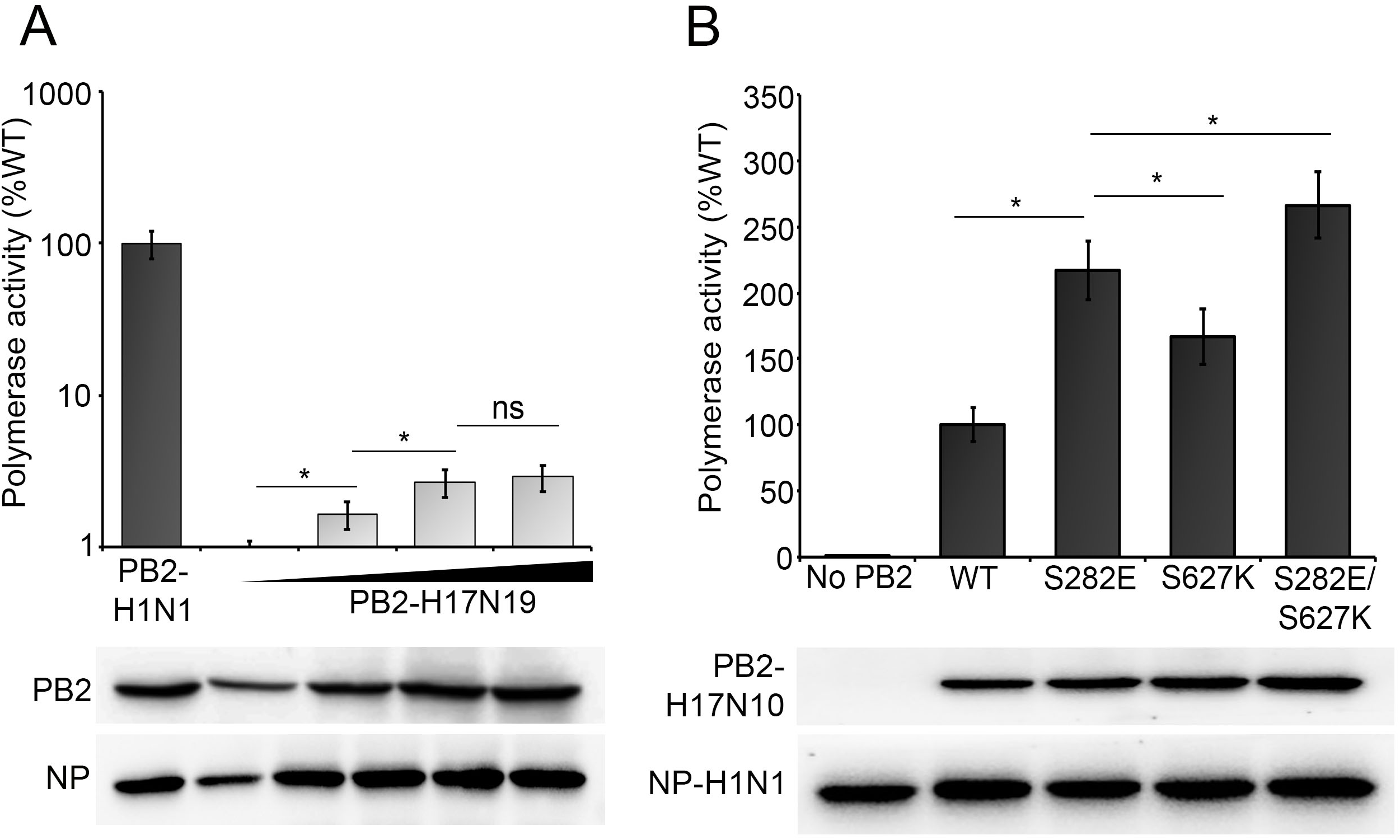
Introduction of S627K and S282E mutations in H17N10 PB2 protein boosts activity of the chimeric polymerase. **A**. Luciferase based reporter assay was performed by reconstituting the Influenza A/H1N1/WSN/1933 RNP in HEK293T, either with H1N1 PB2 or with different amount of H17N10 PB2 expression plasmid. n=3±standard deviation. *p<0.05 one way ANOVA when compared to previous set. **B**. Luciferase based reporter assay was performed by reconstituting the Influenza A/H1N1/WSN/1933 RNP with either wild type H17N10 PB2 or mutant H17N10 PB2 protein. n=3±standard deviation. *p<0.05 one-way ANOVA when compared to PB2 wildtype or S282E or S627K.

residues at 627 and 282 positions with human signature lysine and glutamic acid residues in the H17N10 PB2 protein in order to generate either single mutants (S627K and S282E) or double mutant (S672K/ S282E). Chimeric polymerases and RNPs were reconstituted either with wildtype or mutant H17N10 PB2 proteins to measure their ability to support viral RNA synthesis. Interestingly, substituting of the bat virus specific serine residues with human virus specific glutamic acid or lysine boosted the polymerase activity by 100 percent and 50 percent respectively thereby providing evidence that serine at 282 and 627 position of H17N10 PB2 protein may restrict the activity of bat influenza polymerases in human cells (Fig8.B). Generation of the double mutant S282E/S627K, harboring both of the human virus specific signature residues, showed minor enhancement compared to the single mutants, hence suggesting that these restriction factors in the polymerase does not act in cumulative fashion. Together, our data identified the PB2 S282 as a key residue which in association with PB2 S627 restricts the activity of the bat influenza PB2 protein and hence virus polymerase in human cells.

## Discussion

Higher genetic plasticity of the bat influenza viruses has been shown to facilitate their rapid adaptation in mice (26, 27), hence revealing their potential of cross species transmission in non-bat host species. Interestingly, these adapted bat viruses replicate poorly in ferrets (26, 27) and in human derived cell lines (26), which could be attributed to the suboptimal activity of the viral polymerase in non-bat host species (7). In this work, we have identified a unique molecular signature in the PB2 subunit of bat influenza virus polymerase, which is responsible for its restricted activity in human cell lines. Extensive sequence and structural comparison between different influenza virus polymerases led us to the identification of a previously uncharacterized “S-E-S” motif in the PB2 subunit, which is completely conserved in conventional influenza A viruses, but is replaced with a “S-S-T” motif in bat influenza virus. Alteration of this canonical “S-E-S” motif with bat like “S-S-S” significantly reduced the polymerase activity and hence the fitness of influenza A/H1N1/WSN virus. In corroboration with these results, replacing the bat specific “S-S-T” with semi-canonical “S-E-T” motif results in significant enhancement in the activity of the chimeric polymerase reconstituted with PB1 and PA from conventional IAV and PB2 from bat IAV. Extensive investigation of the molecular mechanism revealed that the complete integrity of the S-E-S motif, specifically the negatively charged glutamic acid at the center of the motif, is critical for supporting RNA synthesis activity of the polymerase and replacing this glutamic acid either with alanine or with the bat specific serine residue severely attenuates the same. Clearly, this S-E-S motif, specifically the 282^nd^ E residues serves as a key molecular determinants of the fitness of classical influenza A virus polymerase in a host independent manner. On the other hand, our work also suggests that the serine at the 282^nd^ position of the bat influenza virus PB2 may serve as inherent restriction element impeding its activity in human cells. It is interesting to note that the bat specific serine residue at the PB2 627^th^ position also imparts similar restriction to the viral polymerase activity in a host independent manner.

The evolutionary history of the bat influenza viruses and their ancestry relationship with other flu variants are not clear (16). However, the new world bat influenza viruses (H17N10 and H18N11 subtypes) have been designated as distinctly divergent entity of IAVs (7, 8). As per the phylogenetic backdating, the new world bat IAVs possibly have separated from all of the conventional lineages around 650 years ago (16). This is supported by the fact that the amino acid sequences of the internal segments of the new world bat viruses tolerate higher diversity in contrast to the conventional IAVs and hence form a distinct outgroup (8). It is possible that the bat influenza viruses served as the ancestor for the all other IAVs, which during the course of adaptation into other host species acquired species specific signatures in different internal gene segments. The change of serine to glutamic acid at 627^th^ (avian) and 282^nd^ positions of PB2 may thus had served as crucial adaptation signatures of the virus from bat to not-bat hosts, which have conferred additional fitness to the viral RNA polymerase in terms of replication in corresponding host species. In another possible scenario, bat and non-bat IAVs may have segregated from a common ancestor (27), where the bat specific serine residues may got selected over glutamic acid in order to generate a slower replication variant of the influenza virus, which might be suitable in terms of establishing host-parasite equilibrium in the bat species. It is interesting that the old-world bat IAVs (H9N2), which shows relatively lesser divergence from the conventional variants (16), harbors glutamic acid at the 627^th^ and 282^nd^ positions of PB2 (28), further supporting the importance of these residues in adaptation of the viruses from bat to non-bat hosts or vice versa.

It is interesting to note that bat influenza virus polymerase, specifically the PB2 subunit, harbors higher number of serine residues in the solvent exposed surface in comparison to the conventional IAVs (table 1,). For example, all of the aspartic acid or glutamic acid residues identified in this study are serine or threonine in bat virus polymerase. It is tempting to speculate that phosphorylation of some of these serine residues, specifically at the 627^th^ and 282^nd^ positions, is important for proper functionality of the polymerase and that is why these residues may get adapted to negatively charged glutamic acid or aspartic acid residues in conventional IAVs, so that the dependence over the host kinases could be bypassed. In opposite sense, a glutamic acid to serine adaptation could also impart functional regulation to the bat virus polymerase, which could be controlled via phosphorylation by host kinases. A comparative phospho-proteome analyses of the of the bat and non-bat influenza virus polymerase in various host species may shade light upon this possibility of such naturally occurring phosphomimetic adaptation sites in the conventional influenza A viruses. From a different perspective, mutation of these serine residues to glutamic acid within the bat influenza virus polymerase may also increase its replication fitness, which together with the adaptive mutations in the HA and NA (26, 27) could lead to the spillover of the bat viruses in non-bat host species.

### Materials and methods

#### Cell Culture

Human embryonic kidney (HEK293T), Madin-Darby Canine Kindey (MDCK), Human lung epithelial carcinoma(A549), *Pteropus alecto* kidney (PaKi) cells were maintained in DMEM (Gibco), supplemented with 10% FBS, 1X Penicillin –Streptomycin solution and 1X GlutaMAX at 37°C temperature, 98% relative humidity, in presence of 5% CO2. DF-1(chicken fibroblast) cells were maintained at 39°C temperature, 98% relative humidity, in the same culture media.

#### Plasmids and viruses

All genes were derived from the influenza A (A/WSN/33) or influenza B (B/Brisbane/60/2008) viruses. Polymerase proteins and NP were expressed in cells from the plasmids, pCDNA3-PB2-FLAG (encoding a C-terminal FLAG tag), pCDNA3-PA, pCDNA3-PB1, pCDNA6.2-NP-V5 (encoding C terminal V5 tag for Influnza A) and pcDNA3.1-NP for Influenza B. The PB2 ORF of Influenza A/H17N10/guatemala/060 was cloned also under the CMV promotor in pCDNA3.1 vector with C terminal 3X FLAG epitope tag. vNA-luc reporter plasmids encodes firefly luciferase in the negative sense flanked by UTRs from the NA gene derived from A/H1N1/WSN/1933 and B/Brisbane/60/2008 cloned under RNA Polymerase I promoter and terminator in pHH21 vector system for expression in HEK293T cells and in pGHH21 for expression in DF1 cells. Viruses were prepared using the pBD bi-directional reverse genetics system using pBD plamids for polymerase subunits and NP and pTMΔ RNP for expression of all other segments.

#### Site directed mutagenesis

Site directed mutagenesis primers were designed by quick change primer design software (Agilent technologies). Mutation was performed by PCR amplification of the plasmids using Pfu Turbo DNA Polymerase enzyme (Agilent technologies) followed by DpnI digestion and transformation in E. coli DH5α cells. Mutations were confirmed by Sanger sequencing.

#### Polymerase Activity Assays

HEK293T or DF-1 cells were reverse transfected in triplicates using Lipofectamine 3000 with plasmids expressing PA, PB1, PB2-FLAG,NP-v5/NP and vNA-Luc. Cells were harvested 36 h.p.t, polymerase activity measured using the Luciferase Assay System (Promega) on GloMax20/20 luminometer (Promega). Equivalent PB2 and NP protein expression were confirmed by western blotting.

#### Rescue of recombinant viruses and plaque assay

Co-cultures of HEK293T and MDBK cells were reverse transfected using Lipofectamine3000 Reagent with virus rescue plasmids pTMΔRNP, pBD_PB2-FLAG143, pBD*_PB1, pBD_PA and pBD_NP. 24 h.p.t, the culture media was replaced with Virus Growth Media (1X DMEM, 1X PenStrep, 4 mM GlutaMAX, 0.2% Bovine Serum Albumin (BSA), 25 mM HEPES buffer and 0.5 μg/ml TPCK Trypsin). Supernatants were harvested 48-72 h.p.t and the viruses subsequently amplified in MDBK cells. Plaque assay was performed in MDCK cells. 0.3million cells were seeded in each well of 12 well cell culture plate day before the assay. Virus stocks were diluted in 10-fold series in virus growth media. Cells were infected by the dilutions for 50 minutes at 37°C with intermittent shaking in each 10 minutes. After 50 minutes the infected cells were overlaid with 1:1 mixture of 2.4% avicel and 2X DMEM (2X DMEM, 1X PenStrep, 4 mM GlutaMAX, 0.4% Bovine Serum Albumin (BSA), 50 mM HEPES buffer and 1.0 μg/ml TPCK Trypsin). Plates were harvested at 72 hours post infection, fixed with 70% ethanol and stained with 1% crystal violet. Plaques were counted from the stained plates and titer was calculated.

#### Multi cycle replication kinetics

A549, MDCK and PAKI cell lines were infected with wild type or mutant Influenza A/H1N1/WSN/1933 viruses at MOI of 0.01 in three biological replicates. Supernatants were harvested at 8,16,24,48 and 72 hours post infection. Viral titer in the supernatant was quantitated by performing plaque assay in MDCK cells.

### RNA Isolation, Reverse Transcription and PCR

Supernatants from virus rescue experiments were harvested, clarified by centrifugation at 12,000 g for 10 minutes, and viral RNA was extracted using TRIzol reagent (Invitrogen). Reverse transcription was performed by M-MLV RT PB2-specific primers. PCR amplification of cDNA was done by Phusion DNA Polymerase with PB2-gene specific primer pair. PCR amplified fragments were used for sanger sequencing.

#### Primer Extension

PB1,PA and wild type or mutant PB2 encoding plasmids with a small 77nucletide long viral RNA with 3’ and 5’ UTR of NP (NP77) expression plasmid were transfected in HEK293T cells. 48 hours post transfection the cells were harvested. Total RNA was isolated by TRIzol reagent (Invitrogen). Primer extension was performed by Superscript III reverse transcriptase enzyme (Thermo) and fluorescence labelled primers as described earlier(25). The reactions were separated in Urea-PAGE and the gel imaged in Bio-Rad chemidoc.

#### Co-immunoprecipitations

HEK293T cells were transfected for RNP reconstitution with expression plasmids encoding NP-v5, PB1, PA and FLAG-tagged WT or mutant PB2, along with vRNA (NA segment). Cells were lysed 48 h.p.t. using Co-IP buffer (50 mM Tris-HCl [pH 7.4], 150 mM NaCl, 1 mM EDTA, 1% NP-40, 1% Na-deoxycholate, 0.1% SDS) supplemented with 1X protease inhibitor cocktail (10 μL of 50X PI in 500 μL of Co-IP buffer), Halt phosphatase inhibitor (10 μL for 500uL of lysis buffer), and incubated at 4°C for 20 minutes on a rocker. Lysates were clarified by centrifuging at 20,000g for 20 minutes at 4°C. Total protein samples were separated from the lysate. Lysates were supplemented with 0.5mg/mL BSA and precleared with 20uL of pre-equilibrated Protein A agarose beads. After preclearing the lysates were incubated overnight with the antibody. Next day, pre-equilibrated protein A magnetic beads were added to the lysate and incubated for 1 hour. Protein-A beads were then recovered using magnetic racks and washed thrice with Co-IP buffer. Finally, sample beads were treated with 30 µL of 5X Laemmli buffer, heated at 98°C for 5 minutes, centrifuged at 10,000g for 10 minutes. Samples were separated by SDS-PAGE and identified by Western blotting.

#### Western blotting

Cell lysates were separated by SDS-PAGE and transferred to methanol-activated PVDF membrane (Bio-Rad) using Trans-Blot Turbo Transfer System. The membrane was blocked in 5% skimmed milk solution in 1X TBST at room temperature for 2 hours. Primary antibody incubation done at 4°C in rocking condition for overnight. Next day, after washing three times with 1X TBST, the membrane was incubated with HRP-conjugated secondary antibody at room temperature for 1 hour. Before developing, the blots were washed thrice with 1X TBST. Chemiluminescent substrate applied over the blot and incubated for 3-5 minutes and developed in BioRad chemi-doc.

#### Immunofluorescence Assay

A549 cells grown on coverslips were transfected with plasmids encoding wild type or mutant PB2 proteins. 24 hours post-transfection, cells were washed in PBS, fixed with 3% formaldehyde (20 minutes at room temperature), quenched with 0.1M Glycine, permeabilized with 0.1% Triton-X 100 in PBS (10 minutes at room temperature). After blocking with 3% BSA (20 minutes at RT), cells were incubated with primary antibody (anti-FLAG) for 1 hour at room temperature and washed thrice. Secondary antibody (Alexa Fluor 488-conjugated donkey anti-mouse IgG) incubation done for 40 minutes at room temperature. DAPI staining was done along with the secondary antibody incubation. After secondary antibody incubation cells were washed thrice with PBS and once with nano-pure water and mounted on slide with Fluoroshield (Sigma Aldrich). Images were taken in fluorescence microscope (Leica).

#### Bioinformatics and structural analysis

Specific PB2 protein sequences were obtained from Influenza Research database (www.fludb.org) by selecting data type-protein, Virus type-A, proteins-PB2 and separately selecting Host-Human(n=36243), Avian(n=19762) and Bat(n=7). The aligned FASTA files were viewed and analyzed in Bioedit sequence alignment editor(29). Multiple sequence alignment was performed in Bioedit software by using ClustalW followed by manual trimming. Logo plots were generated by WebLogo server (weblogo.berkeley.edu/logo.cgi)using the aligned FASTAfiles (http://www.ebi.ac.uk/pdbe/prot_int/pistart.html) (30). PDBePISA tool (31) was used to analyze the solvent assessible surface area of PB2 proteins from the following PBD file-4WSB,4WSA,6RR7 and 5D98(19, 20, 32). Structural alignment of the PB2 protein from trimeric polymerase crystal structure was performed in UCSF Chimera software.

### Statistical analysis and data analysis

Graph preparations and statistical analysis were done in Microsoft Excel software. Densitometric analysis was performed in Bio-Rad Image Lab software.

## Acknowledgement

We sincerely acknowledge Prof. Andrew Mehle (University of Wisconsin Madison) for his comments on the manuscript and providing valuable resources. A.M. thanks DBT, Ramalingaswami re-entry fellowship (BT/RLF/Re-entry/02/2015), SERB, Early Career Research Award (ECR/2017/001896) and MHRD, “Scheme for Transformational and Advanced Research in Science” {STARS/APR2019/BS/369/FS (Project ID: 369)} for financial support. Individual fellowship for SB (File No.09/081(1301)/2017-EMR-I_),NK(File No.09/081(1316)/2017-EMR-I_) and AD(File No.09/081(1405)/2020-EMR-I_) was provided by Council of Industrial and Fundamental research, Govt. of India. We sincerely acknowledge Dr. Gayatri Mukherjee from School of Medical science and Technology, IIT Kharagpur, for providing fluorescence based gel imaging facility.

